# Temperature and Water Quality-Related Patterns in Sediment-Associated *Symbiodinium* Communities Impact Symbiont Uptake and Fitness of Juvenile Acroporid Corals

**DOI:** 10.1101/199802

**Authors:** Kate M. Quigley, Line K. Bay, Bette L. Willis

## Abstract

The majority of corals acquire their photo-endosymbiont *Symbiodinium* from environmental sources anew each generation. Despite the critical role that environmental availability of *Symbiodinium* plays in the potential for corals to acclimate and adapt to changing environments, little is known about the diversity of free-living *Symbiodinium* communities and how variation in these communities influences uptake and *in hospite* communities in juvenile corals. Here we characterize *Symbiodinium* community diversity in sediment samples collected from eight reefs representing latitudinal and cross-shelf variation in water quality and temperature regimes. Sediment-associated *Symbiodinium* communities were then compared to *in hospite* communities acquired by *A. tenuis* and *A. millepora* juveniles following 11 – 145 days of experimental exposure to sediments from each of the reefs. Communities associated with juveniles and sediments differed substantially, with sediments harbouring four times more unique OTUs than juveniles (1125 OTUs vs. 271). Moreover, only 10.6% of these OTUs were shared between juveniles and sediments, indicating selective uptake by acroporid juveniles. The diversity and abundance of *Symbiodinium* types differed among sediment samples from different temperature and water quality environments. *Symbiodinium* communities acquired by juveniles also differed among the sediment treatments, despite juveniles having similar parentage. Moreover, *Symbiodinium* communities displayed different rates of infection, mortality, and photochemical efficiencies. This study demonstrates that the biogeography of free-living *Symbiodinium* types found within sediment reservoirs follows patterns along latitudinal and water quality environmental gradients on the Great Barrier Reef. We also demonstrate a bipartite strategy for *Symbiodinium* uptake by juvenile corals of two horizontally-transmitting acroporid species, whereby uptake is selective within the constraints of environmental availability.

## 1. Introduction

Symbiosis between dinoflagellates in the genus *Symbiodinium* and scleractinian corals is fundamental for the formation, health, and survival of coral reefs. Over 75% of coral species establish this relationship anew each generation through the acquisition of *Symbiodinium* from environmental sources, a process termed horizontal transmission (Baker, 2003; Fadlallah, 1983). Environmental conditions on tropical reefs, including the Great Barrier Reef (GBR), are changing rapidly, with increases in mean temperatures, pCO_2_ concentrations, and nutrients of particular concern (Donner, 2009; Lønborg et al., 2016). As climate change and other human impacts continue to devastate reefs worldwide, knowledge of the capacity of corals to expand or change their symbiotic partners to types better suited to changing climate conditions may help predict species and community survival. Equally as important, knowledge of *Symbiodinium* availability and the environmental conditions that shape the composition of free-living communities is essential for projecting how environmental change will impact symbiont acquisition and coral physiological flexibility.

Environmental pools of *Symbiodinium* are thought to influence the uptake of *Symbiodinium* by both larvae and juvenile corals (Adams et al., 2009; Cumbo et al., 2013; Nitschke et al., 2015), although the full extent of free-living *Symbiodinium* diversity is uncharacterised. Heritability estimates of the *Symbiodinium* community in acroporid juveniles also suggest that *in hospite* communities are partly regulated by symbiont availability in the environmental pool and suggest that this association is flexible and adaptable (Quigley et al., 2017b). For example, communities in the sediments act as a link between *in hospite* communities in adult corals, which continually re-seed the environmental pool, and the establishment of symbiont communities in juveniles through re-seeding of the environmental pool (Nitschke et al., 2015), and have been shown to significantly increase acquisition of *Symbiodinium* in larvae compared to the water column (Adams et al., 2009). However, lack of knowledge of the diversity, distribution and abundance of free-living dinoflagellates is currently limiting our understanding of environmental impacts on the establishment of coral-*Symbiodinium* symbioses.

Sediments, the water column, coral rubble, and algal/cyanobacterial surfaces all have distinct *Symbiodinium* communities in the Pacific and Caribbean oceans (reviewed in Quigley et al. Book Chapter). Comparative studies of Caribbean and Hawaiian reefs demonstrate that the dominant *Symbiodinium* clades found in local corals reflect environmental diversity in the respective region, with clades A and B being the most abundant in both the water column and sediments in the Caribbean, and clades A and C being the most abundant on Pacific reefs (Takabayashi, *et al.*, 2012). Clade A was also recovered from coral larvae exposed to sediments, and B and C from juveniles exposed to reef waters in Japan (Adams et al., 2009). In addition, larvae infected using sediments sourced from the northern and central GBR contained *Symbiodinium* C1, C2, and D, supporting the presence of these types in sediments from these regions (Cumbo et al., 2013). Deep sequencing technology has begun to reveal free-living diversity of *Symbiodinium* (Cunning *et al.*, 2015), but further studies across diverse habitats are needed to fully elucidate biogeographical patterns in the distribution and abundance of these important symbionts.

Physiological responses of *in hospite* versus cultured *Symbiodinium* differ in response to thermal, irradiance and nutrient regimes at the clade and type level (Baker, 2003; van Oppen et al., 2009), but there is limited knowledge of whether variation in these environmental factors shapes free-living distributions of this symbiont. There is some evidence from *in hospite* and cultured studies that temperature regimes may influence local availability of thermally tolerant (clades D and F (Sawall et al., 2014)) versus thermally sensitive *Symbiodinium* types (A and B, although A1 is thermo-tolerant (e.g. Karim *et al.* 2015)). Biogeographic patterns may also follow water quality gradients, as high levels of suspended sediments change light environments and nutrient levels (Storlazzi et al., 2015). Parameters affecting irradiance, such as mud content and Secchi disc depth, correlate well with distribution patterns of *in hospite Symbiodinium* on the GBR (Cooper et al., 2011) and also with the abundance of clade C in some coral species in the Caribbean (Garren et al., 2006). Nitrogen (N) and phosphorus (P) utilization varies amongst *Symbiodinium* clades/types during photosynthesis (e.g. greater nitrate uptake by C versus D (Devlin, 2015; Iluz and Dubinsky, 2015)) and is potentially host-regulated and limiting for *Symbiodinium in hospite* (Rees, 1991). Inorganic forms are readily available in interstitial waters of carbonate sediments (Entsch et al., 1983) and may therefore represent important nutrients in structuring environmental *Symbiodinium* populations and communities. Although environmental availability of different *Symbiodinium* types is essential for establishing symbioses and for the potential formation of more stress-tolerant partnerships, it is unknown if gradients in temperature, light or nutrients influence the distributions of free-living *Symbiodinium*.

To estimate the capacity of corals to respond to climate change through the establishment of symbiosis with novel or locally-adapted symbionts, the diversity and abundance of *Symbiodinium* types within sediment communities were characterized using deep sequencing across latitudinal (temperature) and cross-shelf (water quality) gradients. Aposymbiotic juveniles of *Acropora tenuis* and *A. millepora* were exposed to eight sediment treatments, and subsequent *in hospite* diversity was characterized after 11 – 145 days of exposure (corresponding to ∼2 week to 6 month old juveniles) to determine if the availability of free-living *Symbiodinium* types impacts uptake during early ontogeny. Finally, the initial infection dynamics and longer term physiological impacts of *in hospite Symbiodinium* communities on the health and survival of coral juveniles were also determined. We demonstrate that juveniles are highly selective in the *Symbiodinium* they take up, but are also influenced by the availability of locally abundant types. Symbiont availability and the corresponding variability in symbiont uptake also correlated with variations in growth and survival of juvenile corals.

## 2. Materials and Methods

### 2.1 Experimental design and sample collection

#### 2.1.1 Sediment collections

Sediments were collected from four sites representing an inshore-offshore water-quality gradient in each of the northern and central sectors of the GBR. Samples were collected in October in both 2013 (northern sector) and 2014 (central sector) just prior to the mass coral spawning in each year. Wallace Islets and Wilkie Island (northern sector) and Pandora and Magnetic Island (central sector) are located inshore, where water quality is characterized by relatively high loads of fine particulate matter and nutrients (Figure S1A, Figure S3). Sites at Great Detached and Tydeman reefs (northern sector) and at Rib and Davies reefs (central sector) are offshore and experience fewer terrestrial inputs. As a consequence, they typically have lower exposure to fine particulate matter and nutrient loads. At each site, surface sediments (top 10 cm) were collected at a water depth of 4 m in three replicate one-litre containers. Sediments were maintained in separate bins supplied with 0.1 µM filtered, flow-through seawater under shaded conditions until experimentation (one bin with three litres of sediment per site in 2013; three replicate bins, each with three litres of sediment per site in 2014). An additional three litres of sediment per site at all northern and central reefs were immediately frozen at -20°C for later analysis. In 2013, experiments were run at Orpheus Island Research Station (OIRS), and in 2014, at the National Sea Simulator (Seasim) at the Australian Institute of Marine Sciences (AIMS). Seawater supplied to sediment treatments at OIRS follow ambient temperature conditions on Orpheus Island reefs and was filtered to 0.1 µM (Figure S1B). Seawater supplied to sediment treatments at AIMS was filtered to 0.4 µM and, during the first month of infection, followed a regime representing the average of seawater temperatures at the four central sector reefs for that time of year; seawater temperatures were then maintained at 27.4°C for the grow-out period (Figure S1A, B). In both years, experimental tanks were shaded to approximately 50 µmol photon illumination. To compare *Symbiodinium* communities in sediments pre- and post-experimental sediment treatments, an additional litre of sediment was frozen after field collection, as well as after 145 days of experimental exposure from central sediment locations.

### 2.2 Breeding and husbandry of *Acropora tenuis and Acropora millepora* juveniles

Eggs and sperm from the broadcast spawning corals *Acropora tenuis* and *A. millepora* were isolated from colonies collected from Wilkie Bay and Orpheus Island in 2013, and from Magnetic Island and Trunk Reef in 2014. Spawning, gamete fertilization and larval rearing were performed as described in (Quigley et al., 2016) (see Supplementary Methods for further details about spawning and settlement). For the northern sediment experiments, larvae were settled onto autoclaved terracota tiles and then randomly placed into one of four experimental treatments that corresponded to sediments from one of the four northern reefs (one tank per treatment), at 11 days post-fertilization (*A. tenuis:* n = 6 tiles per tank, providing a total of 4,012 juveniles across the four sediment treatments; *A. millepora*: n = 2 tiles per tank for the Wilkie sediment treatment, 7 for Wallace, 4 for Great Detached, 7 for Tydeman; 903 juveniles overall). For central sediment experiments, juveniles were settled onto plastic 6-well plates and roughly evenly distributed among the four tanks (*A. tenuis*: n= 3-5 plates per tank; *A. millepora:* 1-2 plates per tank). All surviving juveniles from each treatment were sampled following either 35 days of exposure (days of exposure: d.o.e.) to northern sediments, or over 12 time points between 11 to 145 days of exposure to central sediments, and stored in 100% ethanol at -20°C until DNA extraction (Table S1).

### 2.3 Genotyping *Symbiodinium* communities in sediments and within juveniles

#### 2.3.1 Filtration and concentration of sediments

To concentrate *Symbiodinium* cells prior to DNA extraction, frozen sediments were thawed, then filtered with a series of Impact Vibratory Sieves (500, 250, 125, and 63 µM stainless steel mesh) using filtered seawater (0.1µM filtration) and concentrated into 5.5L of 0.1µM filtered seawater. To fully detach *Symbiodinium* cells from sediment grains, the filtration process was repeated five times. All equipment was washed and autoclaved between samples at 124°C and 200kPa for 20 minutes (Sabac T63). The 5.5L of filtrate was concentrated into a pellet (< 50 ml) with serial centrifugation at 4°C: 4500 rcf for 5 min (x2), 10 min (x3), 15 min (x3), 30 min (x1) (Allegra X-15R: Beckman-Coulter). The pellet was stored at -20°C until the time of extraction.

#### 2.3.2 DNA extraction

*Symbiodinium* DNA was extracted from three replicate sediment filtrate samples (each 10 grams) per site (3 x 9 sites, including Orpheus Island), and from pre- and post-experiment samples for central sediments (3 x 4 sites; total n = 39 sediment samples). DNA was extracted using the Mo Bio Powermax Soil DNA Isolation Kit (Carlsbad, CA) following a modification of the manufacturer’s instructions, i.e., including an additional digestion step in a rotating oven with buffer C1 at 65°C and 80 rpm rotation for 30 min. The samples also underwent three, 20-sec, bead-beating steps (6.0 m/s) with 1mm silica spheres (MPBio) on the FastPrep-24 5G (MP Biomedicals). DNA was extracted from 59 and 176 coral juveniles from the northern and central sector sediment studies, respectively, using a SDS digestion and ethanol precipitation protocol (Wilson et al., 2002) (see Table S1 for number of juveniles used per treatment). This extraction included three x 30 seconds of agitation at 4.0 m/s with 1 mm silica spheres at the lysis buffer step.

### 2.4 Sequencing and data analysis

Next generation amplicon sequencing of the ITS-2 locus, data processing, and bioinformatics were performed as detailed in (Quigley et al., 2016, 2017b). Briefly, after quality checks and filtering, a total of 2,188 OTUs were identified based on a custom *Symbiodinium* database. This this total was reduced to 1,562 OTUs retrieved across all juvenile and sediment samples with further quality-checks (further details in Supplementary Methods). Mapped reads were variance-normalized using the ‘*DESeq2*’ package (v. 1.6.3, Love *et al.*, 2014) implemented in R (R Team, 2013). Abundances of *Symbiodinium* OTUs were compared among treatments using negative binomial generalized linear models using ‘*DESeq2*’. Adjusted p-values were derived using the Benjamini-Hochberg Multiple-inference correction for alpha = 0.05. Differential abundances calculated in ‘*DESeq2*’ were expressed in multiplicative (log2 fold) terms within or among treatments (Love *et al.*, 2014).

Seven analyses were performed to compare *Symbiodinium* communities among the different treatments and samples (temporal sampling design in Table S1) using differential abundance testing implemented in ‘*DESeq2*’. Briefly, comparisons included differential abundance testing among sample types (all juveniles vs. all sediment samples), cross-shelf positions (inshore vs. offshore), sectors (northern vs. central sector), and interactions among these factors (type*shore*sector), as well as further comparisons within each of the factors (further details in Supplementary Methods).

Nonmetric multidimensional scaling (NMDS) was performed on variance-normalized OTU abundances using the packages ‘*Phyloseq*’, ‘*vegan*’ and ‘*ellipse*’ with a Bray-Curtis dissimilarity matrix (McMurdie and Holmes, 2013; Murdoch et al., 2007; Oksanen et al., 2013). NMDS analysis does not assume linear relationships between underlying variables, and distances between samples are indicative of their similarity in OTU diversity and abundance (Ramette, 2007). Permutational multivariate analysis of variance was used to determine if *Symbiodinium* communities differed significantly between the factors listed in detail above using the ‘*adonis*’ function in ‘*vegan*’. Correlation coefficients of (variance-normalized, averaged) abundances of each of the 1,562 OTUs across the eight treatments for juveniles and eight sites for sediment samples were calculated and visualized using the package ‘*corrplot*’ (Wei, 2013).

### 2.5 Juvenile physiological measures

#### 2.5.1 Time to infection and survival

The trait “time to initial infection” was measured as the first day that *Symbiodinium* cells were visible throughout the whole juvenile (i.e., within the oral disc, tentacles, and polyp column). Survival was defined as the total number of days an individual juvenile was observed alive, which was determined with a stereo dissection microscope at each sampling date (Table S1). To determine the effect of sediment source on the number of days to infection and the number of days surviving, generalized linear mixed models were fit in ‘*lme4*’ (Bates et al., 2014) (further details about model parameterization in Supplementary Methods).

#### 2.5.2 Photosynthetic measurements

Photophysiological performance of *in hospite Symbiodinium* was assessed in *A. tenuis* juveniles with Imaging Pulse Amplitude Modulated (iPAM) fluorometery and its affiliated software (Walz, Effeltrich, Germany). The iPAM allows resolution at 100 µM, thus provides accurate photosynthetic measures of small juvenile corals (Hill and Ulstrup, 2005). Symbiont densities were large enough to obtain detectable measurements at 41 d.o.e. The actinic light was calibrated with an Apogee quantum sensor (Model MQ-200, UT, USA) with the following settings: measuring intensity = 3, saturation pulse intensity = 3, gain = 2. Juveniles were dark adapted for 10 minutes prior to measurements beginning at 10:00 AM, and maximum potential quantum yield (ratio of maximum to variable fluorescence: F_v_/F_m_) was calculated. This measurement is routinely used to determine photosynthetic productivity in studies of plant physiology (Maxwell & Johnson 2000), reflects the efficiency of PSII (Krause and Weis 1991), and is a widely accepted indicator of stress in corals (Jones et al. 1999).

To detect differences in F_v_/F_m_ among *A. tenuis* juveniles exposed to the four central sector sediment treatments, Generalized Additive Mixed Models (GAMMs) were used to account for non-linear trends over time (Wood, 2006) using the package ‘*mgcv*’ (Wood, 2006, 2008). Penalized regression spline smoothing functions were applied to the interaction between sampling date and site, whilst site itself was also included as a fixed effect. Plate was treated as a random effect and the variance structure was allowed to vary through time using the varIdent weights argument. Temporal autocorrelation from sequential measurements was dealt with using first-order autoregressive correlation structure at the deepest level (plate) (Pinheiro and Bates, 2006). Model selection was performed with AIC and the log-likelihood ratio tests using the ‘anova’ function from the ‘*nlme*’ package. A log-normal distribution was used, as F_v_/F_m_ values are non-integer values inherently greater than zero. ACF plots and normalized residual plots versus fitted values conformed to assumptions of no autocorrelation and heterogeneity of variance.

To examine if there were any relationships between F_v_/F_m_ measurements and normalized OTU abundances in *A. tenuis* juveniles exposed to sediments from central sector sites from day 41 to day 90, nine of the overall most abundant OTUs (OTU2_B1, OTU3_C1, OTU2129_C15, OTU115_C, OTU1_A3, OTU9_A13, OTU4_D1, OTU7_A3, OTU427_C15), which represented the majority of reads retrieved from juveniles from these time points, were selected for correlation analysis of normalized abundances and F_v_/F_m_ values. OTU abundances and F_v_/F_m_ values were averaged across replicate *A. tenuis* juveniles, thereby providing one Spearman’s Rho Correlation coefficient (R^2^) per OTU per time point per sediment site (R base package ‘Stats’ (R Team, 2013)). This metric is robust to non-normal bivariate distributions (Becker et al., 1988), as was the case here.

#### 2.5.3 Environmental covariates

Mud and carbonate content, average sea surface temperatures (SST) from 2000-2006, and 10 water quality measures were used to identify environmental drivers of *exhospite* (sediment) *Symbiodinium* diversity and allow for comparisons with *in hospite Symbiodinium* diversity (Cooper et al. 2011). Water quality, irradiance and temperature data were collected from datasets generated by the Australian Institute of Marine Sciences, the Great Barrier Reef Marine Park Authority, and Department of Primary Industry and Fisheries from 1976-2006, and retrieved from eAtlas (Brodie et al., 2007; De’ath and Fabricius, 2008; De’ath, 2007; Furnas, 2003; Furnas et al., 2005). Values for each covariate per site were extracted from interpolated modelled data using the R packages ‘*dismo*‘ (Hijmans et al., 2013) and ‘*raster*’ (Hijmans), and figures for each were created using the package ‘*mapping*’ and a custom Queensland spatialPolygons file created by Murray Logan (Logan, 2016). Mud and carbonate content were categorized as per Maxwell’s (1968) scheme for the Great Barrier Reef (full description in Supplementary Materials). Water quality measures consisted of: total dissolved nutrients (dissolved inorganic nitrogen (DIN: NO_2_, NO_3_, NH_4_), total dissolved phosphorus (TDP), total dissolved nitrogen (TDN), particulate phosphorus (PP), particulate nitrogen (PN)) and irradiance (suspended solids (SS), chlorophyll *a*, secchi depth). Central sediment sites were further characterized by nutrient profiles and particle size distributions (Supplementary Methods).

To address the highly correlated nature of many of the irradiance and water quality measures, each of the 10 measures were z-score standardized using the scale function in base R, and then summed for each site to create a Water Quality Index measure (WQI), as per methods in Cooper et al. (2011). Correlations between environmental covariates (WQI, mud and carbonate content, SST) and specific *Symbiodinium* OTUs found in the sediments were based on pre-experimental sediment samples (n = 27 samples in total, i.e., northern sediments = 3 reps x 4 sites, central samples = 3 reps x 5 sites; 1160 of the 1562 OTUs). Sixteen types with known taxonomic information (Guiry and Guiry, 2016) of the possible 1160 OTUs from the pre-experimental samples were selected for Generalized Additive Model analysis (GAMS) across 7 of the 9 clades. GAMS and partial plots were constructed using the packages ‘*mgcv*’ (Wood, 2000, 2006, 2008) of normalized abundance data generated from the full data set. Non-significant covariates and regression spline smoothing functions (smoothers) were dropped to achieve final models and normalized abundances of OTUs identified to the same type were summed. A non-metric multidimensional scaling plot (NMDS) of the full data set was constructed using the Bray-Curtis distance metric, with the positioning of environmental covariates as vectors created using the packages ‘*vegan*’, and ‘*ggplot2*’(Wickham, 2009).

## 3.Results

### 3.1 Overall comparison of *Symbiodinium* communities between coral juveniles and sediments across all factors and time points

*Symbiodinium* communities differed significantly between juvenile corals and sediments when data were combined across all samples of sediments, juveniles (both species) and time points (Figures 1, 2, Table 1). Of the 1,562 OTUs that passed the E-value filter, 72% (1,125) were found only in sediment samples, whereas 17.3% (271) of OTUs were unique to juveniles. Only 10.6% (166) were shared between juveniles and sediments (Figure S2C). *In hospite* and *ex hospite* diversities were only moderately correlated among sites (mean R^2^= 0.229 ± 0.04, Figure 3).

**Table 1.** Summary of Permutational Multivariate Analysis of Variance tests to compare *Symbiodinium* community composition among juveniles and sediments. Factors are defined as follows: type (all juveniles or all sediment samples), shore (inshore or offshore) and sector (northern or central).

| Factor | Df | F | R<sup>2</sup> | P |
| --- | --- | --- | --- | --- |
| Type | 1 | 20.841 | 0.057 | 0.001 |
| Shore | 1 | 15.371 | 0.425 | 0.001 |
| Sector | 1 | 37.308 | 0.103 | 0.001 |
| Type*Shore | 1 | 5.226 | 0.014 | 0.001 |
| Type*Sector | 1 | 9.217 | 0.025 | 0.001 |
| Shore*Sector | 1 | 4.490 | 0.012 | 0.002 |
| Type*Shore*Sector | 1 | 2.610 | 0.007 | 0.004 |
| Total | 273 |  | 1.0 |  |

**Table 2.** Spearman’s rho R^2^ correlation coefficients calculated for each of the nine most highly abundant OTUs in juveniles compared to F_m_ values per site. Although A13 (OTU9) was one of the most abundant types in juveniles overall, this OTU was only highly abundant time points 1-3. Therefore, no correlation coefficients were calculated due to the zero abundance of this particular A13 OTU (OTU9) by e point 4 when F_v_/F_m_ measurements commenced.

| Clade/Category | OTU | Type | Offshore |  |  |  | Inshore |  |  |  |
| --- | --- | --- | --- | --- | --- | --- | --- | --- | --- | --- |
|  |  |  | Davies |  | Rib |  | Magnetic |  | Pandora |  |
| | | | $F_v/F_m$ | $R^2$ | $F_v/F_m$ | $R^2$ | $F_v/F_m$ | $R^2$ | $F_v/F_m$ | $R^2$ |
| B | 2 | ↑*S. minitum* | ↑ | 0.41 | ↑ | 0.2 | ↑ | **0.8** | ↓ | -0.2 |
| C | 3 | ↑C1 | ↓ | -0.08 | ↑ | **0.8** | ↑ | 0.4 | ↑ | 0.2 |
| C | 2129 | ↑C15 | ↓ | -0.01 | ↑ | **0.8** | NA | NA | ↓ | -0.4 |
| C | 115 | ↑C | ↓ | -0.23 | ↑ | 0.2 | ↑ | **0.8** | ↓ | **-0.8** |
| A | 1 | ↑A3 | ↓ | **-0.74** | ↓ | **-0.6** | ↑ | **1.0** | ↓ | **-1.0** |
| A | 9 | ↑A13 | NA | NA | NA | NA | NA | NA | NA | NA |
| D | 4 | ↑D1 | ↑ | 0.059 | ↓ | **-1.0** | ↑ | 0.4 | ↑ | 0.4 |
| A | 7 | ↑A3.2 | ↑ | 0.24 | ↓ | -0.25 | ↑ | 0.1 | ↑ | 0.4 |
| C | 427 | ↑C15 | ↑ | 0.21 | ↑ | 0.4 | ↑ | 0.4 | ↓ | -0.4 |

**Figure 1.**
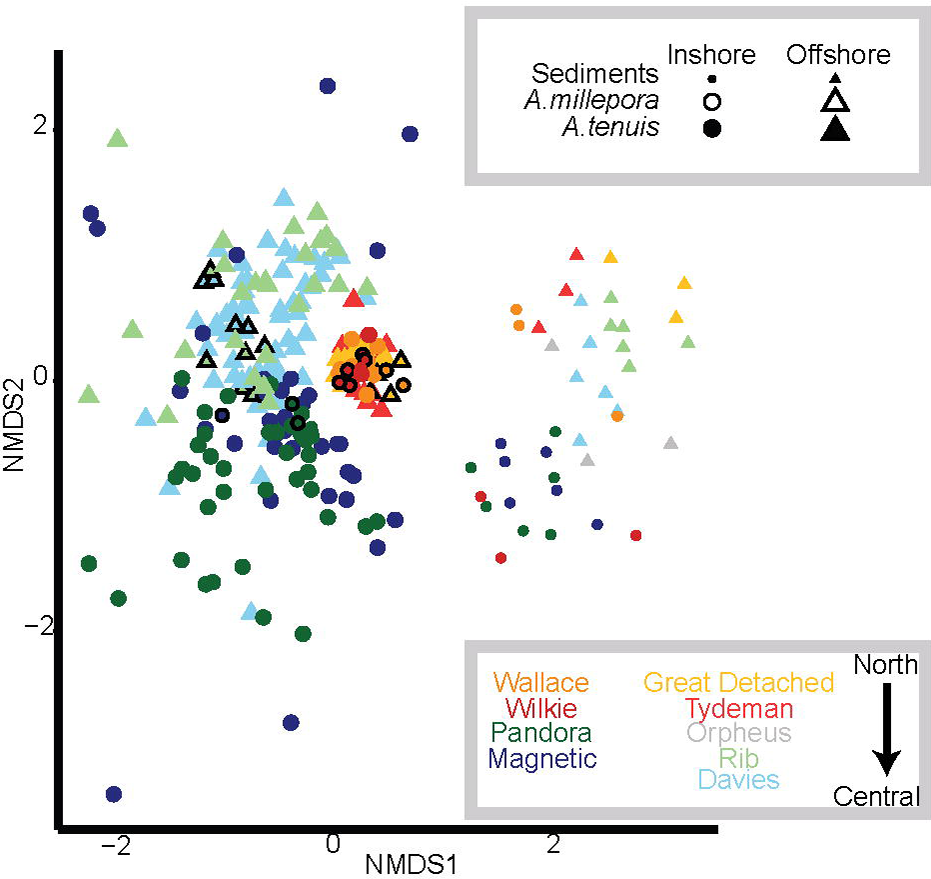
Nonmetric multidimensional scaling (NMDS) of variance-normalized abundances of *Symbiodinium* OTU’s using Bray-Curtis distances. Symbols and their sizes represent: sediment samples (small circles and triangles), *A. millepora* juveniles (black outlined symbols) and *A. tenuis* juveniles (no outlines). Colours denote sector and cross shelf origin of sediment samples, where yellow and red denote northern sediments and juveniles exposed to them; blue, green and grey colours denote central sediments and juveniles exposed to them. Darker shades represent inshore reef sediments and treatments (Wallace, Wilkie, Pandora, Magnetic); lighter shades represent offshore reef sediments and treatments (Great Detached, Tydeman, Orpheus, Rib, Davies).

**Figure 2.**
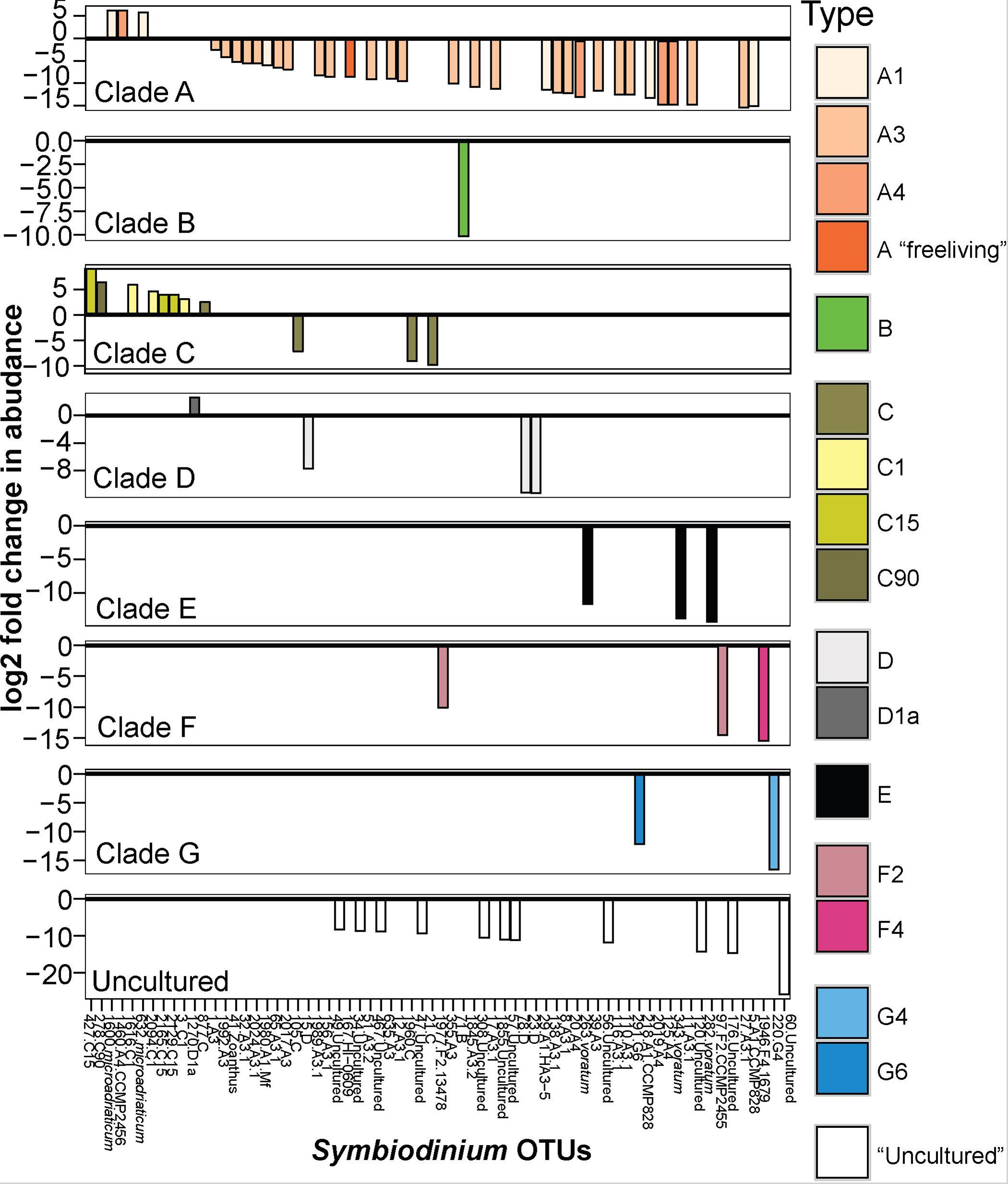
Significant log2 fold changes in normalized abundances of *Symbiodinium* OTUs in sediment-exposed juveniles compared to sediment samples. Colours represent the classification of different *Symbiodinium* types. Positive values above the y = 0 line (in bold and black) denote *Symbiodinium* OTUs found in significantly greater abundances in juveniles compared to sediments. Note the different scales on the y-axis.

**Figure 3.**
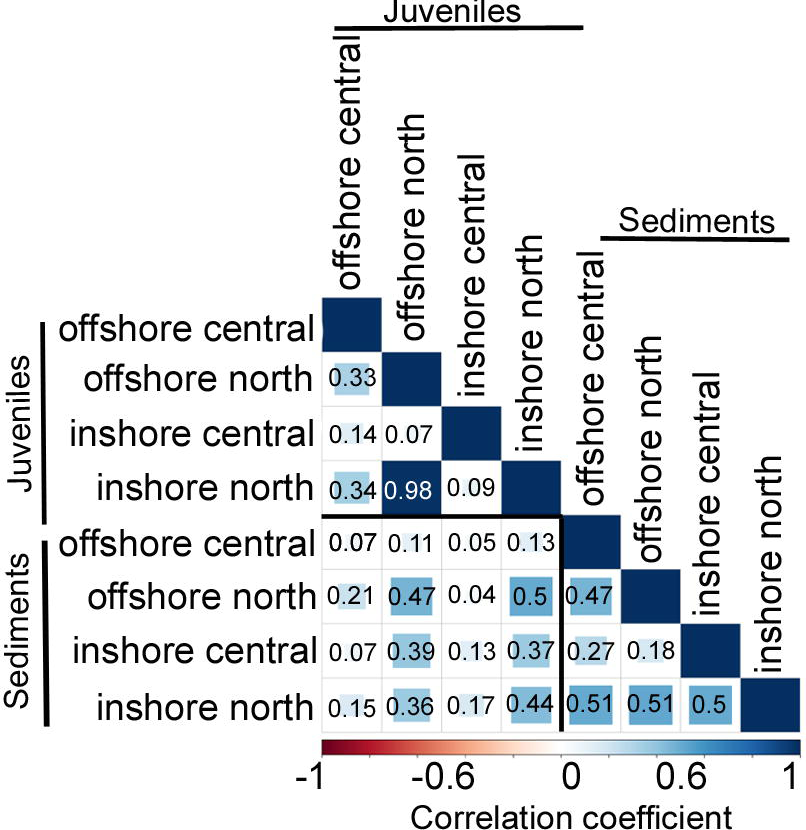
Matrix of pairwise correlation coefficients between *Symbiodinium* communities associated with juveniles versus in the sediments across latitudes (northern vs. central GBR) and water quality gradients (inshore vs. offshore). The scale bar represents coefficients from -1 in red (100% negatively correlated) to 1 in dark blue (100% positively correlated).

OTUs shared between juveniles and sediments belonged predominantly to clades A and C (39 and 27 OTUs, respectively). *Symbiodinium* in clades D - H were also present in both sample types, but at lower diversities (< 40 OTUs in total) (Figure S2). Overall, within-clade diversity was much lower in juveniles (< 100 OTUs) than in sediments (< 300 OTUs). The abundance of 69 OTUs differed significantly between sediment and juvenile samples and included types from clades A - G (Figure 2, B-H adjusted p-values, log2 fold adjusted p-values according to Benjamini-Hochberg (B-H) multiple-inference correction for alpha < 0.05). In general, juveniles had up to 2.3 times greater abundances of an A4 and an A1 species (*S. microadriaticum*), as well as of multiple types belonging to C1, C15, C90, and D1a, compared to the sediments (Figure 2). In contrast, the sediments had greater abundances of types within clades B, E, F, G and “uncultured” *Symbiodinium* that have not yet been identified taxonomically. The sediments also held a larger abundance of diverse types within clade A, including free-living types from A1, A3, and A4.

### 3.2 Distribution and abundance of *Symbiodinium* communities within sediments

#### 3.2.1. Overall patterns in sediment communities across all locations

Clades A and C were the most abundant *Symbiodinium* clades in the sediments (A: 19.5% of total sediment reads, 177 of 1291 OTU sediment diversity, C: 11.5%, 148 of 1291, respectively), particularly a free-living type from clade A (OTU_10). A majority of the unique OTU diversity retrieved from sediment samples was identified as “uncultured” *Symbiodinium* (i.e. novel *Symbiodinium* without taxonomic descriptions, >300 OTUs), which represented 82% of all uncultured *Symbiodinium* OTUs across the whole dataset (i.e., sediment and juvenile samples combined; Figure S2). Approximately 100 unique OTUs were represented from each of clades A, C, D, E, and F, along with a smaller number from clades B, I and G (Figure S2). Greater than 91% of all clade E OTUs were retrieved exclusively from the sediments. *Ex hospite Symbiodinium* community diversity in sediment samples was only moderately correlated among sites (mean R^2^ = 0.41 ± 0.06, Figure 3, Supplementary Results), mostly due to strong differences in diversity across the inshore to offshore gradient.

#### 3.2.2 Inshore versus offshore sediment community comparisons

*Symbiodinium* communities in the sediments differed significantly between inshore and offshore locations (*p=*0.001, Table 1; Figure 1). In total, the abundances of 58 OTUs differed between inshore and offshore sediments (Figure 4). Inshore sediments had up to 3.3 times greater abundances of D1a, D1, a B type, and F3 compared to offshore sediments (B-H adjusted p-values < 0.05, Figure 4). Offshore locations had up to 3.6 times greater abundances of types from clades A (including A1, A3, A4), *S. voratum* (clade E), and F2.

**Figure 4.**
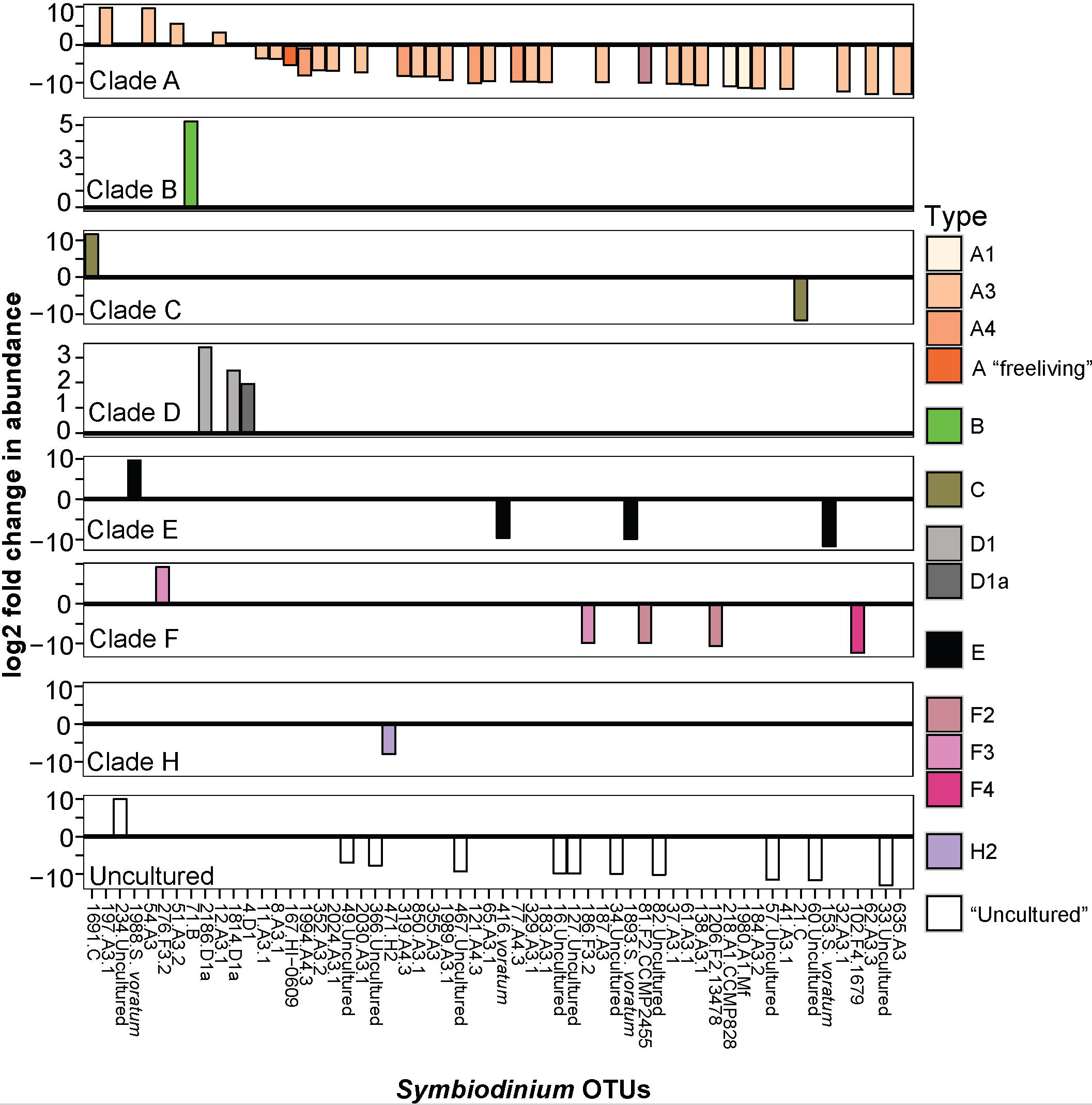
Significant log2 fold changes in normalized abundances of *Symbiodinium* OTUs in inshore compared to offshore sediments. Positive values above the y = 0 line (in bold and black) denote Symbiodinium OTUs found in significantly greater abundances in inshore sediments. Note different scales on the y-axis.

#### 3.2.3 Northern versus central sector sediment community comparisons

*Symbiodinium* communities within northern sector sediments also differed significantly from those detected within central sector sediments (*p=*0.001, Table 1; Figure 1). The abundances of 47 OTUs differed significantly between northern and central sector sediments (Figure 5). Northern sediments had up to 3.6 times greater abundances of 10 different A-types, including A1, A3, A13, and up to 3.1 times greater abundances of C and C1 types, which were not found in sediments from central reefs (B-H adjusted p-values < 0.05, Figure 5). There were also up to 3.6 times higher abundances of F3, F5, G6, and *S. voratum* in northern sediments. Central sediments had up to 2.6 times greater abundances of a B type and a new A type (Oku17), as well as 3.1 times greater abundances of cultured A *Symbiodinium* (HI-0609).

**Figure 5.**
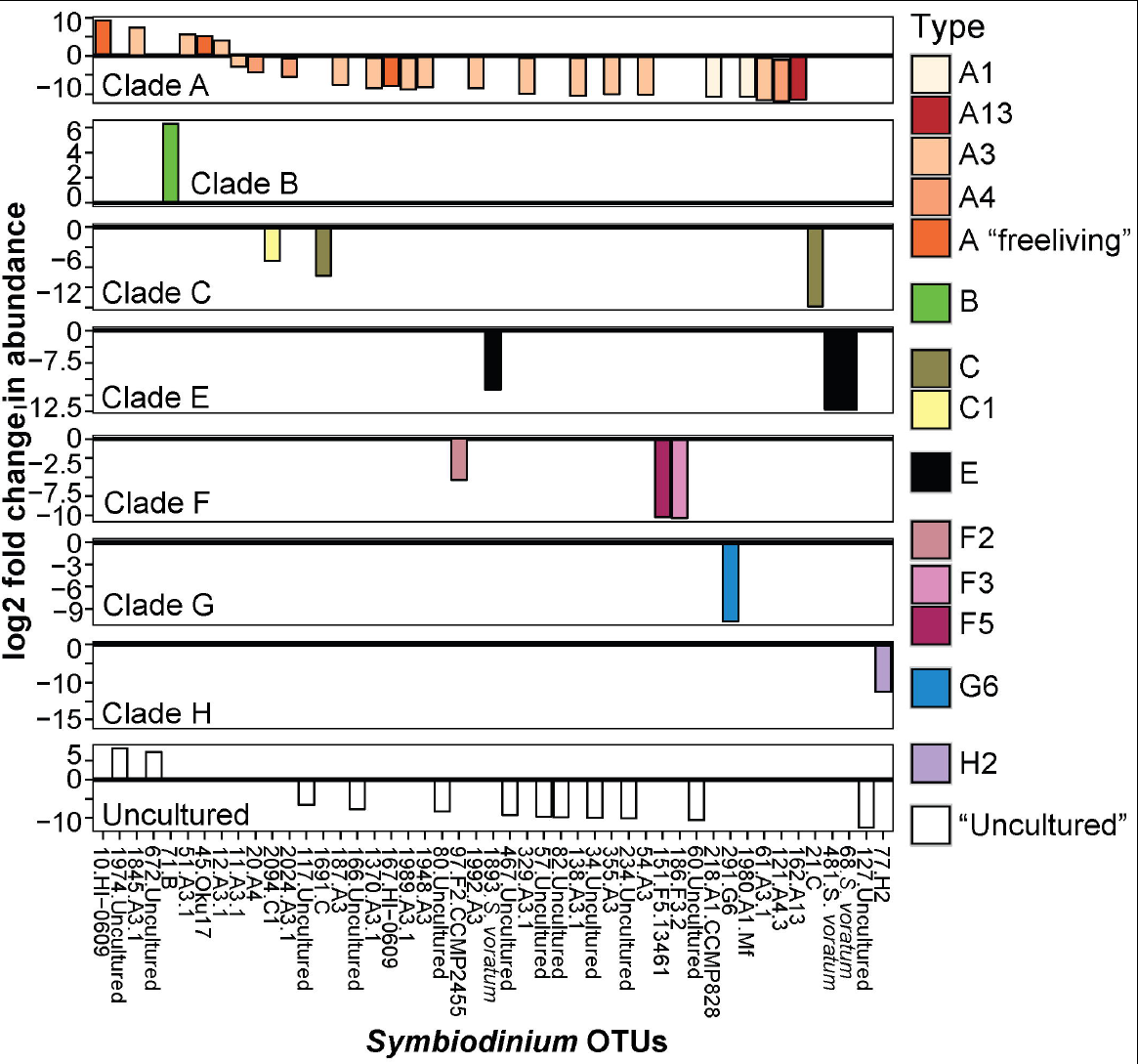
Significant log2 fold changes in normalized abundances of *Symbiodinium* OTUs in central compared to northern sector sediments. Positive values above the y = 0 line (in bold and black) denote *Symbiodinium* OTUs found in significantly greater abundances in central sediments.

### 3.3 Distribution and abundance of *Symbiodinium* communities within juveniles (27-41 d.o.e.)

#### *3.3.1 Symbiodinium* communities in juveniles compared among sediment treatments

Of the OTUs unique to juveniles, clade C had the greatest type diversity (99 OTUs), although each type was relatively rare in abundance, thus C types comprised only 9.8% of juvenile reads overall. In terms of abundance, unique juvenile OTUs belonged predominantly to B1, with this type representing 79.5% of all reads unique to juveniles (Figure S2). As found for sediments, *in hospite Symbiodinium* community diversity was only moderately correlated among juveniles exposed to sediments from the different sites (mean R^2^ = 0.33 ± 0.14, Figure 3, Supplementary Results). In addition, patterns in community composition for juveniles exposed to sediments from inshore versus offshore sites differed between sectors (*p=*0.004, Table 1).

#### 3.3.2 Inshore versus offshore sediment treatments

The early *Symbiodinium* community in juveniles differed significantly when juveniles were exposed to inshore versus offshore sediments (*p=*0.001, Table 1; Figure 1). The abundances of 12 OTUs differed significantly in juveniles exposed to inshore versus offshore sediments (Figure 6A). Juveniles exposed to inshore sediments were dominated by B1 and had up to 3.2 times greater abundances of D1, D1a, and A4 compared to juveniles exposed to offshore sediments (B-H adjusted p-values < 0.05). Juveniles exposed to offshore sediments were dominated by A3, D1, and C15, and had up to 2.9 times greater abundances of A3 and C15 types compared to juveniles exposed to inshore sediments (B-H adjusted p-values < 0.05).

**Figure 6.**
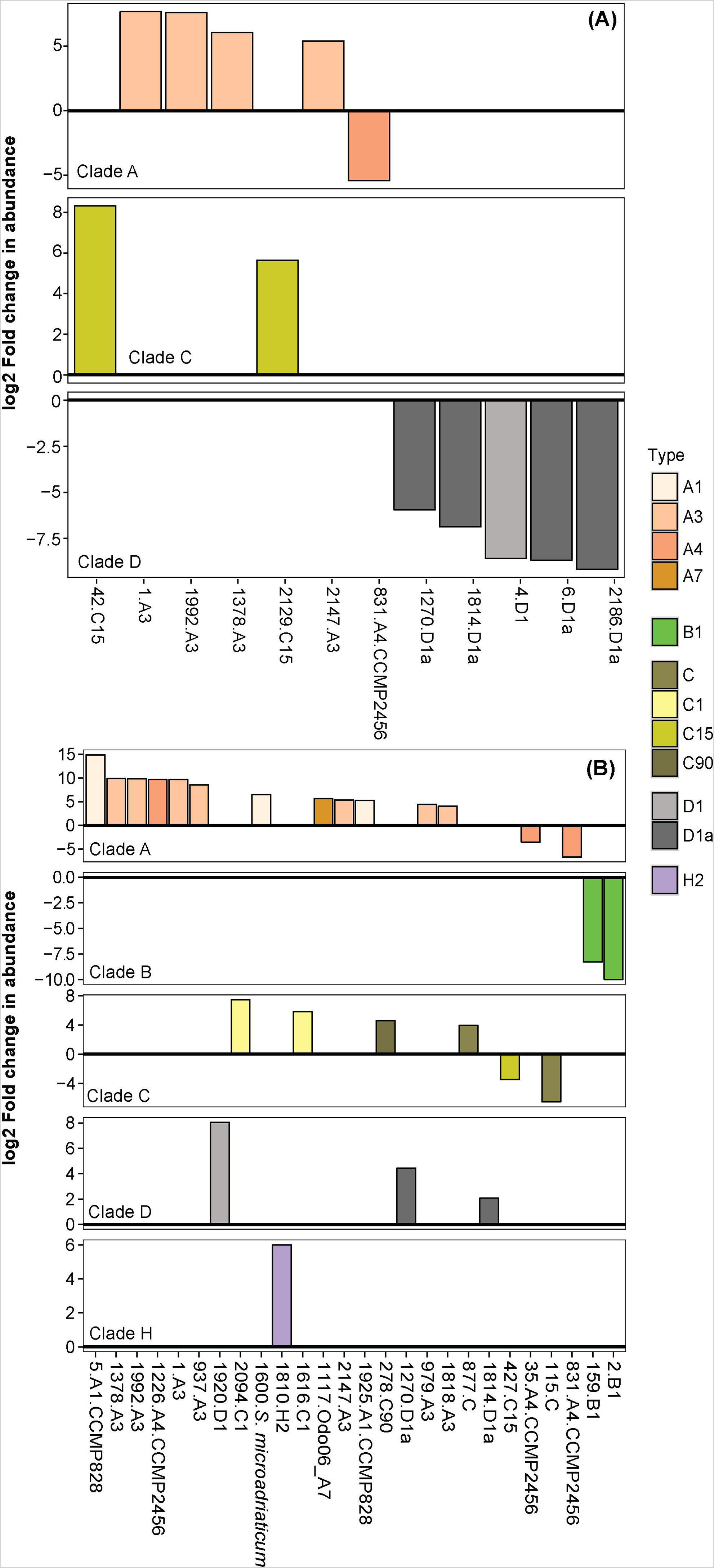
Significant log2 fold changes in normalized abundances of *Symbiodinium* OTUs in juveniles exposed to: **A)** inshore versus offshore sediments, and **B)** central versus northern sector sediments. OTUs above the y = 0 line (in bold and black) are *Symbiodinium* OTUs that were found in significantly greater abundances in: **A**) offshore than in inshore sediments, and **B**) in northern sector sediments than central sediments.

#### 3.3.3 Northern versus central sector sediment treatments

The early *Symbiodinium* community in juveniles exposed to northern versus central sector sediments differed significantly in the abundances of 26 OTUs (*p=*0.001, Table 1; Figure 6B). Juveniles exposed to northern sediments were dominated by A3 and D1 and had up to 3.9 times greater abundances of A1, A3, A7, and A4, D1a, D1, C1, C90, and C (B-H adjusted p-values < 0.05). Juveniles exposed to sediments from the central sector were dominated by B1, C15, C1, and C-types, and had up to 3.3 times more B1, C, C15, and A4 compared to juveniles exposed to northern sediments.

### 3.4 Temporal variation in *Symbiodinium* communities among juveniles exposed to central sector sediments

*Symbiodinium* communities in *A. tenuis* juveniles exposed to central sector sediments differed significantly over time (permutational multivariate analysis of variance: Df _6,140_, F = 2.71, R^2^ = 0.09, *p* = 0.001), by reef (Df _3,140_, F = 7.86, R^2^ = 0.13, *p* = 0.001), and over time in a reef-dependent manner (reef^*^time point, Df _18,140_, F = 1.24, R^2^ = 0.13, *p* = 0.017). At the earliest time point sampled (day 11), juveniles exposed to sediments were all dominated by A3.2 (Davies, Magnetic and Rib sediments) and A13 (Pandora) (Figure 7). Subsequently, types C1, D1, C and A3 varied in abundance by site and over time (B-H adjusted p-values < 0.05, Table S3). In particular, large community shifts were seen 8 days later (day 19) at each site, with communities in juveniles exposed to central offshore reef (Davies and Rib) sediments more closely resembling each other, whereas communities in juveniles exposed to central inshore reef (Magnetic and Pandora) sediments remained distinct. At this second time point, the dominant types (A3.2 or A13) were replaced by B1 (Pandora sediment treatment), D1 (Magnetic), C15 (Davies) and C15/A3 (Rib). By the fifth time point (day 48), *Symbiodinium* communities in juveniles exposed to offshore sediments were beginning to resemble each other, particularly in their abundances of C15, A3 and C1. In contrast, it wasn’t until day 75 that *Symbiodinium* communities in juveniles exposed to inshore sediments started to resemble each other, most notably in their abundances of B1, C1 and D1. After approximately three months of exposure to sediments (day 90), the abundances of C1 and C15 had increased to become dominant in juveniles exposed to sediments from all sites, following steady increases between days 41 and 90 (Figure 7, B - H adjusted p-values < 0.05, Table S3).

**Figure 7.**
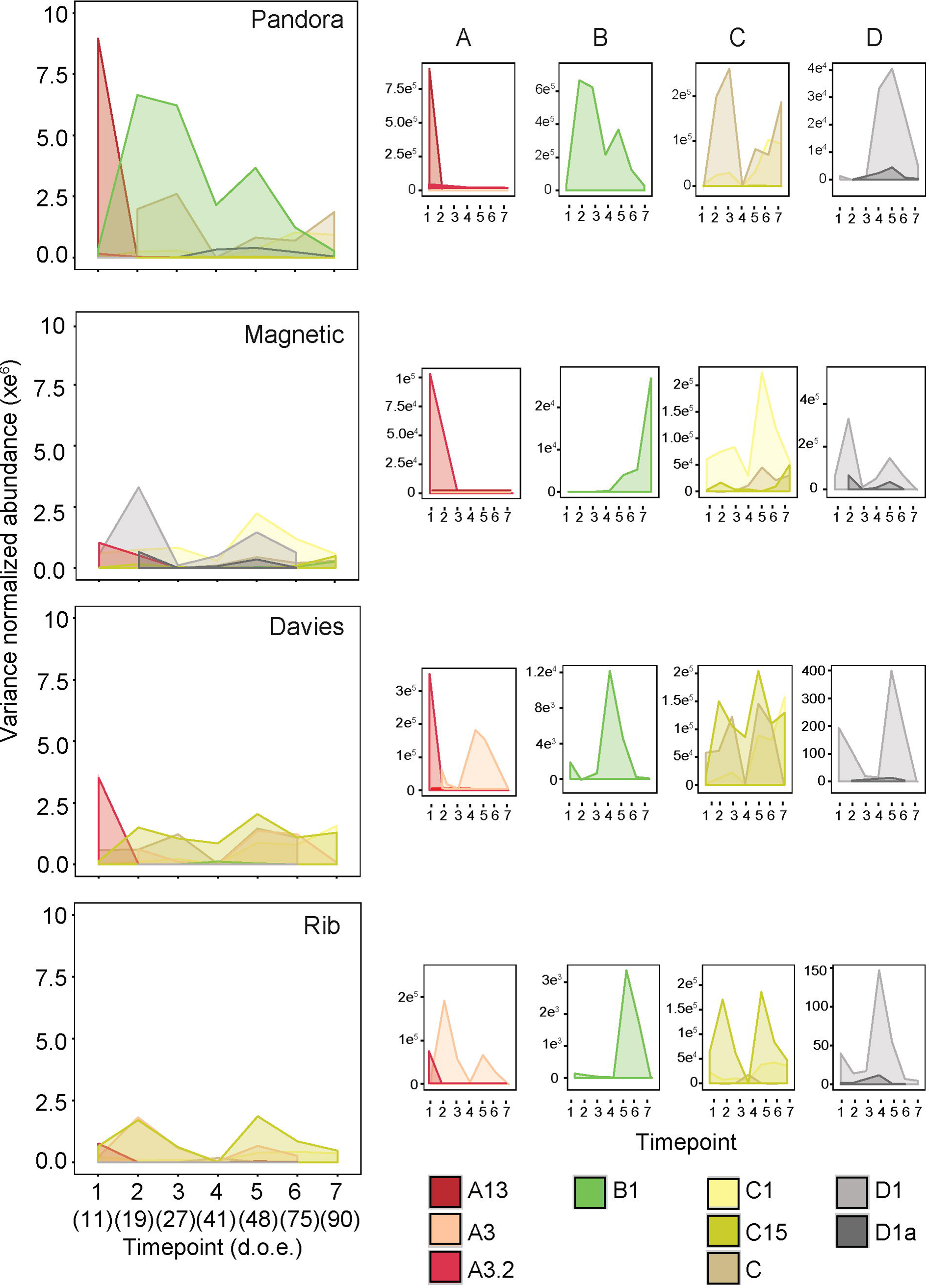
Population dynamics of key *Symbiodinium* types in A. tenuis juveniles exposed to central sector sediments. Abundances of *Symbiodinium* types were variance normalized.Analyses were restricted to 7 time points between day 11 and day 90 of sediment exposure (d.o.e.). The larger panels are all to the same scale, whilst the smaller panels for each site have y-axes that are scaled to best show abundance dynamics for each *Symbiodinium* type. Colours represent nine key Symbiodinium types, whose abundances differed significantly among time points, as well as among sites.

### 3.5 Time to infection and survival of juveniles exposed to central sector sediments

Infection was significantly more rapid in juveniles exposed to offshore sediments (19.1 days ± 0.09) than in juveniles exposed to inshore sediments (24.3 days ±2.13) (negative binomial generalized linear mixed model: *p =* 0.014), with no effect of location (*p =* 0.45) (Figure 8A). The cross-shelf (shore) effect was predominantly driven by significantly lower mean times to infection for juveniles exposed to offshore Rib and Davies sediments (19±0 days and 19.3±0.15 days, respectively) compared to inshore Pandora sediments (26.2 days ±4.37, *p =* 0.0145). Juveniles exposed to inshore Magnetic sediments had intermediate times to infection (22.54 days ± 1.06) that were not significant once the fixed effect of gradient was dropped.

**Figure 8.**
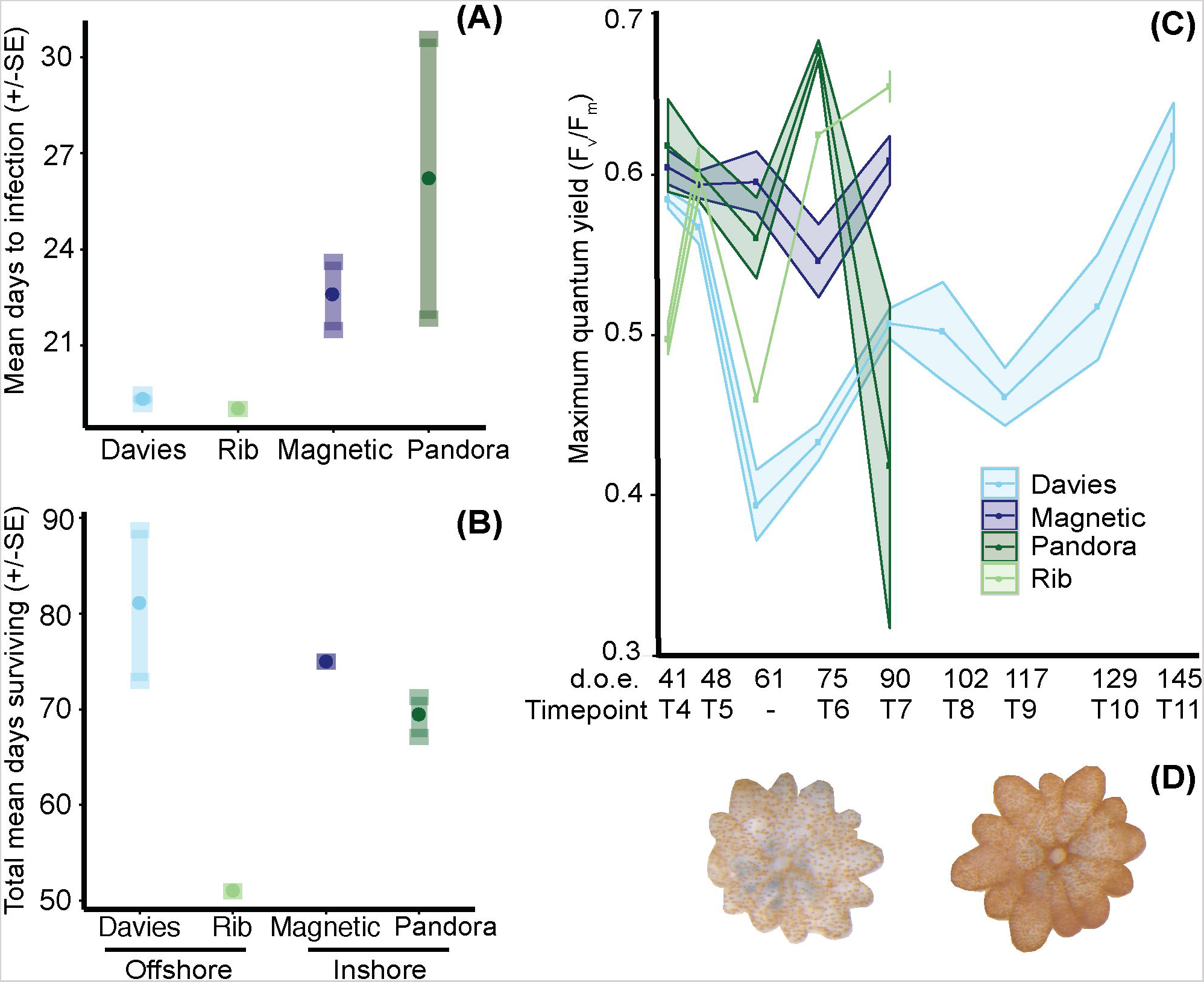
Time to infection and survival of *Acropora tenuis* juveniles exposed to sediments from central GBR sites. Inshore reefs are in darker shades and offshore sites in lighter shades (see x-axis). **A**) Mean time to infection ± SE per sediment treatment.Circles represent means; minimum and maximum values are located at the end of each standard error whisker. **B**) Juvenile survival in days (mean ± SE) per sediment treatment. **C**) Maximum quantum yield (F_v_/F_m_) of juveniles from 52 to 166 days post-fertilization. Due to low survivorship and cumulative sampling of Rib recruits, only one individual was measured at days 59 and 72, and two individuals at day 102, which represents the number of days of exposure to treatments. **D)** Two juveniles from the Davies sediment treatment at day 117, highlighting the level of variation in symbiont proliferation within juveniles exposed to sediments from different sites.

Survival of juveniles did not differ when juveniles were exposed to offshore (68.9±5.7 days) versus inshore (71.95±1.2) sediments (Poisson GLMM: *p =* 0.179), although significant differences did exist at the site level (*p =* 0.002) (Figure 8B). Juveniles exposed to Rib sediments survived significantly fewer days (51 ± 0 days) compared to those exposed to Davies (80.8 ± 7.9, TPH: *p =* 0.003) and Magnetic (75 ± 0, TPH: *p =* 0.03) sediments. However, survival did not differ significantly between juveniles exposed to Rib versus Pandora sediments (69.1 ± 2.1, TPH: *p* = 0.1). Survival of juveniles exposed to sediments from Davies, Magnetic and Pandora did not differ significantly (TPH: *p =* 0.5 - 0.9).

### 3.6 Photo-physiology of juveniles exposed to sediment treatments

#### 3.6.1 Temporal patterns in photochemical efficiency compared among juveniles exposed to central sector sediments

The photochemical efficiency (F_v_/F_m_) of juveniles varied significantly among sediment treatments and over time (Figure 8C; GAM: *p =* 0.0283, Table S7). F_v_/F_m_ values for juveniles exposed to Davies sediments were initially ∼0.6, but dropped sharply by day 48, and then generally increased until day 145. In comparison to juveniles exposed to Davies sediments, temporal patterns in F_v_/F_m_ values differed significantly for juveniles exposed to sediments from the other three locations: Magnetic (t=2.275, *p =* 0.0236), Pandora (t = 2.137, *p =* 0.0334), and Rib (t = 1.997, *p* = 0.0467). Differences in F_v_/F_m_ yields among juveniles exposed to sediments from different locations were also correlated with distinct shifts in *Symbiodinium* communities in these juveniles, for example, the increase in A3 coincided with decreases in F_v_/F_m_ (Figures 7, 8C; Supplementary Results).

### 3.7 Variation in *Symbiodinium* communities between sediment-exposed juveniles of *A. tenuis* and *A*. *millepora*

In northern sediment treatments, juveniles of both *A. millepora* and *A. tenuis* were dominated by *Symbiodinium* A3 and D1, and to a lesser extent by D1a in *A. millepora.* Overall, *Symbiodinium* communities did not differ between *A. tenuis* and *A. millepora* juveniles exposed to northern sediments after 35 days (permutational multivariate analysis of variance: Df _1,58_, F = 2.036, R^2^ = 0.034, *p* = 0.095), although the abundance of nine OTUs varied between species (B-H adjusted p-values < 0.05, Table S4). In contrast, overall *Symbiodinium* communities hosted by juveniles of these two species did differ significantly after 27 – 30 days of exposure to central sediments (Df 1,41, F = 4.83, R^2^ = 0.107, *p* = 0.001). *A. tenuis* juveniles exposed to central sediments were more diverse and had significantly greater abundances of six A types, B1, and C1 (B-H adjusted p-values < 0.05, Table S5) compared to *A. millepora*, which was characterized by additional diversity within C15, D1 and D1a types.

### 3.8 Environmental covariates of *Symbiodinium* communities in sediments

WQI, SST, carbonate and mud content correlated with the normalized abundances of *Symbiodinium* types from clades A, B, C and D, but not types within clades E, F and G (Figure 9, Figure S5, Table S7). Responses of free-living *Symbiodinium* to these environmental covariates were generally linear, in contrast to the non-linear responses of *in hospite Symbiodinium* found in an earlier study (Cooper et al., 2011). Types from clade A were most variable in the covariates that influenced their distributions, with *S. natans* increasing in abundance with increasing WQI and decreasing SST, and decreasing in abundance with lower carbonate and mud content. This contrasted with A3 abundances, which decreased with increasing WQI. The abundance of type A2 was significantly influenced by SST, with greater abundances at higher temperatures. B2 and B4 increased in abundance with increasing SST and sand content (low mud). C1 increased in abundance as carbonate content decreased, whereas none of the environmental covariates were particularly important in explaining the distributions of types C15, C3 or C90. Interestingly, temperature did not significantly explain D1 or D1a abundance, which significantly increased as carbonate decreased in contrast to *in hospite* patterns (i.e., higher abundance in offshore locations Figure 6a). *S. voratum* (E) distributions increased with less carbonate. Finally, none of the environmental covariates tested here explained patterns in the abundances of types *S. kawaguti* (F1), G3 and G6, suggesting that other, unmeasured environmental covariates are important in explaining the distributions of these types. Correspondingly, nutrient profiles and percent of sediments in the < 2000 µM particle size class differed significantly among locations (Supplementary Results, Figure S4A and B, Table S6).

**Figure 9.**
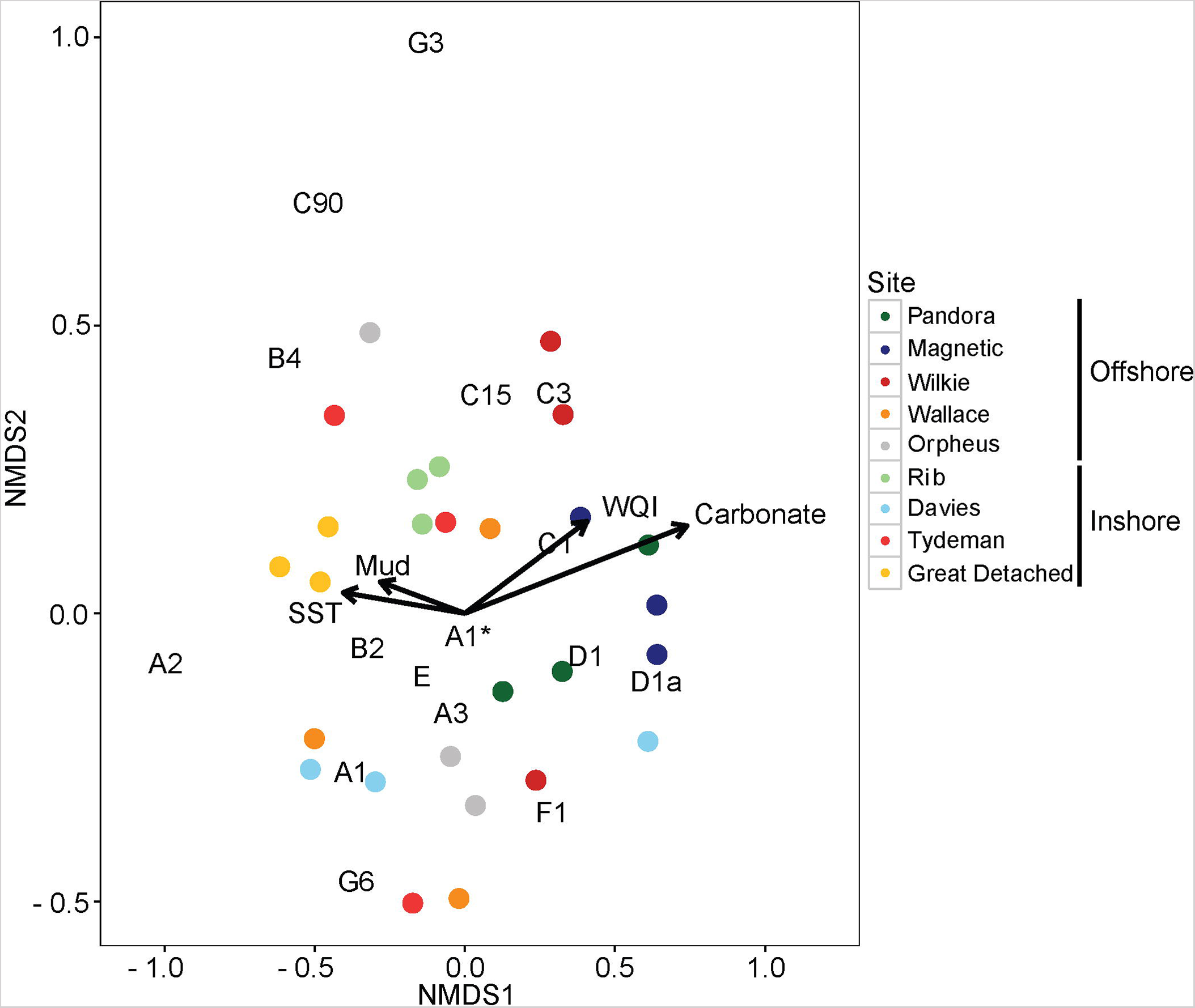
Nonmetric multidimensional scaling (NMDS) of variance-normalized *Symbiodinium* type abundances (summed OTUs that belong to the same type) in pre-experimental sediment samples using Bray-Curtis distances. Northern sediments are in yellow and red, and central sediments are in grey, blue and green colours. Inshore and offshore sediment samples are in dark and light hues, respectively. Environmental covariates (mud, carbonate, Sea Surface Temperature, and Water Quality Index) are overlaid as vectors. Large vector values for mud and carbonate represent low percent mud and carbonate (see Materials and Methods). *Symbiodinium* types in black are also overlaid onto the ordination space. As an example, high normalized abundances of types D1 and D1a correspond to high mud content (i.e. low vector values of mud). A1^*^ is equivalent to *S. natans*.

## 4. Discussion

Comparisons between free-living, sediment-associated *Symbiodinium* communities and endosymbiotic communities in juveniles of the horizontally-transmitting corals *Acropora tenuis* and *A. millepora* reveal a bipartite strategy governing *Symbiodinium* acquisition. Low overlap between OTUs in the sediments and those taken up by juveniles indicates that corals are capable of a degree of selectivity in the symbionts they acquire from environmental sources. At the same time, a component of the variation in *Symbiodinium* communities acquired by juveniles reflected variation amongst sediment sources, indicating that local environmental availability also plays a role, albeit a lesser one, in structuring *in hospite Symbiodinium* communities. Variation in the diversity and abundance of *Symbiodinium* types in sediment sources across a thermal gradient between northern and central sector sites and across inshore-offshore water quality gradients in each sector suggests that environmental conditions influence biogeographical partitioning of free-living *Symbiodinium* communities. Detection of previously unknown levels of *Symbiodinium* diversity in the sediments, combined with evidence that the composition of *in hospite Symbiodinium* communities influences infection rates, survival, and photo-physiology of coral juveniles, has implications for local adaptation of both free-living *Symbiodinium* populations and symbiotic partnerships.

### 4.1 *Symbiodinium* uptake by *Acropora* juveniles is selective but varies with local availability

Only 10.6% of the 1,562 *Symbiodinium* OTUs recovered in this dataset were shared between juveniles and sediments, and OTUs taken up did not necessarily reflect those dominant in sediments. In combination, these findings do not support previous suggestions that the acquisition of *Symbiodinium* from environmental sources is “promiscuous” in early life-history stages of corals (Cumbo et al., 2013). Instead, the low overlap found between these two communities is consistent with a degree of selectivity in juvenile uptake, a conclusion supported by other studies that have compared seawater or benthic communities with those *in hospite,* in association with either adult hard corals (Huang et al., 2013; Pochon et al., 2010; Sweet, 2013) or juveniles of both octocorals and acroporids (Coffroth et al., 2006; Yamashita et al., 2013). Despite only a small proportion of *Symbiodinium* types being taken up out of the wide range of environmental *Symbiodinium* available, *Symbiodinium* communities in juveniles were not highly conserved amongst locations (with the exception of inshore/offshore northern juveniles). Such differences in *in hospite* communities among sediment source treatments suggest that local availability of *Symbiodinium* in the sediments can influence *in hospite* juvenile diversity to a degree. These results align with previous estimates of the influence that environmental availability of *Symbiodinium* has on the formation of those communities in *A. tenuis* juveniles (σ^2^_E_ = 71%, (Quigley et al., 2017b). For example, juveniles exposed to inshore sediments, which had higher abundances of clades B and D than offshore sediments, concomitantly had higher abundances of these two clades than juveniles exposed to offshore sediments. Although *in hospite* communities did not reflect a majority of the *Symbiodinium* types in sediments, linkages between local sediment diversity and *in hospite* communities for some symbiont types may have important implications for local adaptation of coral hosts.

### 4. 2 Predisposition for up-take of specific *Symbiodinium* types

Although local availability of free-living *Symbiodinium* was a contributing factor to the structure of *in hospite* symbiont communities, overall, correlation analyses revealed low similarity in the abundance and diversity of *Symbiodinium* between juvenile and sediment communities at the type/OTU level. The a general lack of congruence between sediment communities and communities found in juveniles exposed to them suggests that there is also a genetic component to symbiont regulation in corals (Quigley et al., 2017a, 2017b). A predisposition for acquiring certain *Symbiodinium* types is supported by the acquisition of clade C symbionts in offshore sediment treatments when their comparative availability in offshore sediments was low, potentially signifying the presence of recognition mechanisms, consistent with findings that clade C ultimately dominates adult communities of these corals (Abrego *et al.*, 2009). A bipartite system, involving both genetic and environmental contributions to the structuring of symbiont communities, has also been observed in closely-related *Nematostella-*bacterial partnerships (Mortzfeld et al., 2015). Knowledge of the environmental availability of symbionts, in conjunction with an understanding of the genetic mechanisms regulating symbiont uptake by the host, is therefore required to predict the potential for *Symbiodinium* communities in acroporid juveniles to affect coral health through differential selection of symbionts in response to environmental change.

The *in hospite Symbiodinium* communities in juveniles also varied through time, although it is unclear whether temporal patterns reflected ontogenetic changes associated with juvenile development or dynamics among *Symbiodinium* types within juveniles. Similar fine-scale successional patterns over short time-scales (1-2 weeks) in bacterial communities have been tied to developmental (including immunological) remodelling in *Nematostella*, amphibians and insects (Mortzfeld et al., 2015). Lack of change in relevant symbiont availability within the sediments from the start to the end of the experiment indicates that lack of symbionts available for infection was not the underlying reason, but does suggest that active selection by the host or competition among symbionts was driving temporal changes in community structure. Although developmental changes associated with metamorphosis in other invertebrate systems may be more extreme than changes associated with growth in *A. tenuis,* skeletogenesis and developmental changes specific to corals have large impacts on gene expression, and differentiate the juvenile stage from larval and adult expression patterns (Reyes-Bermudez et al., 2009). Accordingly, temporal differences and successional patterns in *Symbiodinium* communities associated with *A. tenuis* juveniles may be tied to changing requirements associated with the onset of specific physiological processes (e.g., calcification) or changing colony morphology (e.g., the onset of vertical colony growth) (Abrego *et al.*, 2009). In either case, a preference for increasing abundances of clade C over clade D symbionts may reflect a need for greater translocation of photosynthates, as provided by clade C in experimental studies (Cantin et al., 2009), which can cause 3-fold greater juvenile growth rates in field studies (Little et al., 2004). Further work is needed to determine if selection for *Symbiodinium* from clade C is tied to calcification or other physiological processes during early ontogeny.

Differences in the *Symbiodinium* communities acquired by juveniles of *A. millepora* and *A. tenuis* highlight the diversity in the evolutionary trajectories that underpin the development of symbioses in these two congeneric species. Although there was limited overlap between *Symbiodinium* diversity in juveniles and the sediments, other coral species may utilize and potentially benefit from the *Symbiodinium* diversity in sediments not taken up by these Acroporid species. Additionally, *A. millepora* juveniles in both the northern and central sediment treatments had order of magnitude greater abundances of *Symbiodinium* D1 and D1a (generally heat tolerant types), than *A. tenuis*. If species-specific preferences for D types exist, then by not hosting thermally tolerant types, *A. tenuis* juveniles may be at a comparative disadvantage under future climate change scenarios. Further studies comparing the selection of *Symbiodinium* across different species are required to fully understand the adaptive benefits of flexibility during symbiont acquisition in juvenile corals.

### 4.3 Differences in diversity among sediment *Symbiodinium* communities reflect variation in water quality, temperature, and sediment characteristics

Four times more unique *Symbiodinium* OTUs were recovered from the sediments than from juveniles, representing 72% (1,125) of the overall OTU diversity detected, and indicating that diversity of the free-living community is high. Whilst results of this study are consistent with previous reports of clades A and C occurring in high abundances in Pacific sediment communities (Manning and Gates, 2008; Takabayashi, *et al.*, 2012), the majority of sediment reads detected here belonged to the “uncultured” category. This indicates that many undiscovered and presently uncharacterized types exist within GBR sediments and may represent a reservoir of diversity with potentially adaptive benefits for corals. The *Symbiodinium* types that were unique to the sediments differed among the four regions (north inshore, north offshore, central inshore, central offshore), with the most extreme differences in type diversity being between the northern offshore and central inshore sediments (R^2^=0.18). Inshore central sediments harboured the most distinct *Symbiodinium* communities found and warrants further exploration.

Differences in the abundances of free-living *Symbiodinium* types were also explained by different environmental covariates. For example, C types dominated communities in cooler, nutrient-poor reefs (central offshore sediments) and A types dominated in warmer, higher nutrient, turbid waters (northern inshore sediments). These same patterns were found in the Arabian Gulf, where C1 dominates *P. verrucosa* colonies in colder, nutrient poor waters, whilst A1 dominates warmer, higher nutrient and more turbid waters (Sawall et al., 2014). Offshore reef corals in Palau were also found to be dominated by C types, whilst warmer inshore corals harboured D1a or D1-4 (Russell et al., 2016). Strong biogeographic patterns of *in hospite Symbiodinium* types associated with a soft coral have also been found within clades (Howells et al., 2009) and may be related to variation in local temperature, nutrient and irradiance regimes. Detection of C types in central offshore sediments and A types in northern inshore sediments in both free-living and *in hospite* communities provides compelling evidence that environmental factors contribute to the structure of both symbiotic and environmental communities.

The diversification and partitioning of *Symbiodinium* communities in sediments as a consequence of exposure to different environmental conditions (e.g., different temperatures, light and nutrients) can be ascribed to balancing selection exerted by selective uptake of lower abundant types by the coral meta-population, which continually inoculates the sediments, maintaining overall high diversity of *Symbiodinium* in sediment reservoirs. This reservoir of diversity within sediments might be important to counter loss of *in hospite* diversity from disturbance events and competitive exclusion amongst symbiont types, as has been observed in diverse soil bacterial communities (Martorell et al., 2015). Recent evidence that environmental conditions explain significant variation in the genetic structure of clade C communities (Davies *et al.*, 2016) also highlights the substantial potential for local adaptation of symbionts. Further work on *Symbiodinium* communities in the sediments is needed to describe this important niche and the potentially stress-tolerant species it may contain, including the sequencing of samples collected across environmental gradients representing more extreme environments.

### 4.4 Changes in diversity and abundance of dominant and background *Symbiodinium* types impact coral juvenile physiology

Variation in juvenile survival amongst sediment treatments in the current study may have been driven by differences in the composition and dynamics of the *Symbiodinium* communities acquired. Even fine-scale changes in the relative abundances of some types may translate to large scale impacts at the colony level (Cunning et al., 2015a; Quigley et al., 2016). For example, small shifts in bacterial community composition in *Nematostella* resulted in changes to host fitness, potentially caused by variation in the immune responses elicited by different symbionts under different selective pressures (Mortzfeld et al., 2015). Alternatively, the early acquisition of certain types may have led to greater long-term juvenile survival. For example, 6-fold greater abundances of C and D1 in juveniles exposed to Davies and Magnetic sediments at the first time point corresponded to higher survival compared to juveniles exposed to sediments from other locations. Early acquisition of C and D1 by *Acropora yongei* juveniles has also been correlated with increased juvenile survival on Japanese reefs (Suzuki et al., 2013). Further work is needed to understand how the dynamics of mixed *Symbiodinium* communities contribute to long-term coral juvenile survival, and how this relationship is modulated by the environment and changing environmental regimes.

Variation in the *in hospite Symbiodinium* community among juveniles may also explain the differences we found in time to infection and photosynthetic parameters among communities. *Symbiodinium* infection/proliferation occurred more quickly when juveniles were exposed to offshore sediments (characterised by more abundant C15, A3 and C-types) compared to inshore sediments (characterised by B1, D-types or C1/C-types). Slower infection by cultured D-types compared to C1 is commonly observed in the lab (K. Quigley, pers. obs.), potentially explaining the slower rates of infection when juveniles were exposed to inshore sediments. More rapid rates of infection in juveniles exposed to Rib and Davies (central offshore) sediment treatments may reflect an increased capacity of A3, C15 and C to infect and proliferate within coral juveniles. The presence of A3, both when dominant or at background levels of abundance, also strongly impacted F_v_/F_m_ in juveniles and provides corroborative support that numerically rare background symbionts contribute to changes in the photo-physiological performance of their hosts (Erwin et al., 2012; Hoadley et al., 2015; Karim et al., 2015; Mortzfeld et al., 2015). Lower F_v_/F_m_ values in juveniles that acquired symbionts from offshore sediments and spikes in chlorophyll fluorescence may indicate that the symbionts acquired were adapted to higher light environments (due to increased sediment particle size or decreased turbidity) and a stress response (Shapiro et al., 2016), respectively. These results highlight that the establishment of symbiosis is a highly dynamic process governed by a range of interacting factors.

### 4.5 Conclusion

Low overlap between *Symbiodinium* communities in the sediments and *in hospite Symbiodinium* communities in juveniles exposed to those sediments suggests that overall, symbiont uptake is a selective process. Nevertheless, local environmental availability of *Symbiodinium* also contributes to *in hospite* community structure, albeit to a lesser degree. Evidence that sediment *Symbiodinium* communities vary across temperature and water quality gradients highlights the potential of changing environments to disrupt environmental sources of *Symbiodinium* diversity and therefore the establishment of symbiosis, an important consideration given that the vast majority of coral species on the GBR take up *Symbiodinium* from environmental sources anew each generation. Newly discovered *Symbiodinium* diversity in coral reef sediments presented here highlights the presence of a hitherto unknown reservoir of diversity that may enable corals to adapt to environmental change through the utilization of symbionts at initially low background abundances or by potentially forming new stress-resistant partnerships. The low overlap in diversity between sediment and *in hospite* biomes, paired with evidence of strong biogeographical partitioning of free-living *Symbiodinium* communities in the sediments, has implications for local adaptation of symbiotic partnerships and highlights the need for studies of how climate change and other anthropogenic stressors might disrupt linkages between free-living and *in hospite Symbiodinium* communities.

## Acknowledgements

We thank Orpheus Island Research Station and the Seasim staff for assistance in the set-up of the filtration and tank systems as well as Marie Strader for *Acropora millepora* juveniles. Assistance with sediment collections was provided by Ray Berkelmans, Melissa Rocker, David Stump and Carly Kenkel; Stephan Boyle provided assistance with the nutrient analysis; Andrew Negri for use of sonication filter; Kathryn Evans for her help with filtering sediments. Permits for coral collections in 2013 were as follows: *A. tenuis* colonies from Wilkie Bay (Permit number: G12/35236.1) and eight *A. tenuis* and six *A. millepora* from south Orpheus Island (Permit number: G13/36318.1). Permits for coral collections in 2014 were as follows: four *A. tenuis* colonies from Magnetic Island (Permit Number: G13/36318.1) and eight *A. millepora* colonies from Trunk Reef (Permit Number: G12/35236.1).

## Authors and Contributors

KQ, LB, BW conceived and designed the experiment. KQ performed the laboratory work, data analysis and figure preparation and wrote the manuscript. KQ, LB, and BW critically revised the draft and approved the final version and are in agreement to be accountable for all aspects of the work.

